## Supplementary figures and images for "Multi-omic brain and behavioral correlates of cell-free fetal DNA methylation in macaque maternal obesity models"

### Supplementary Figure 1

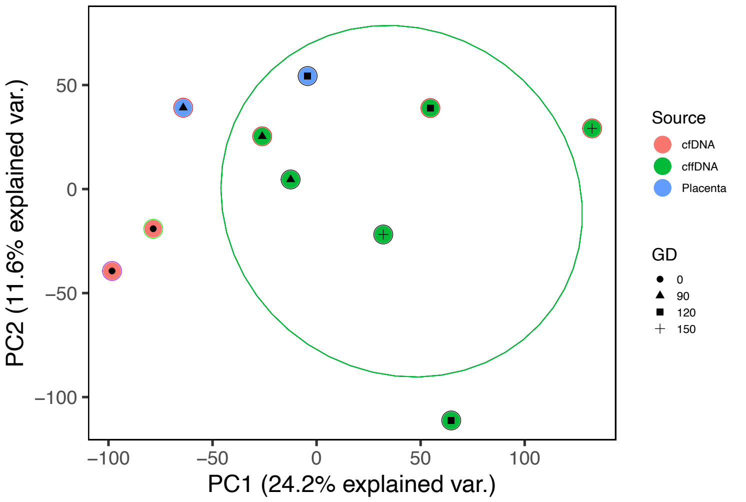

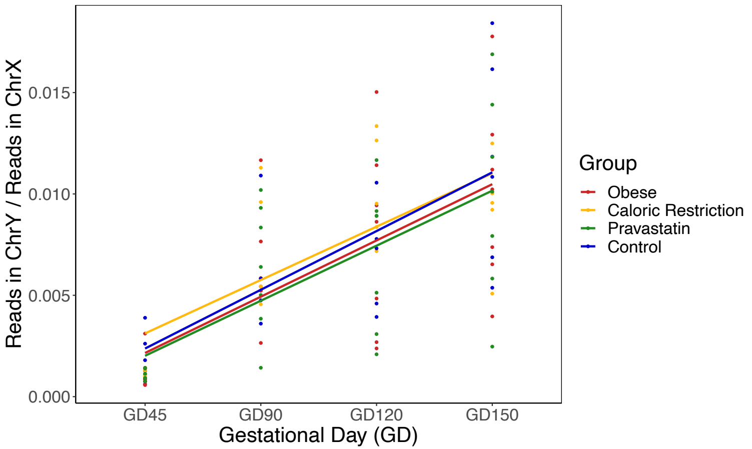

### Supplementary Figure 2

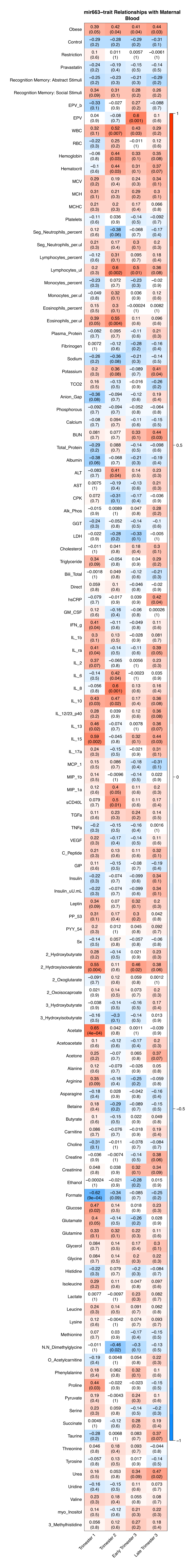

### Supplementary Figure 4

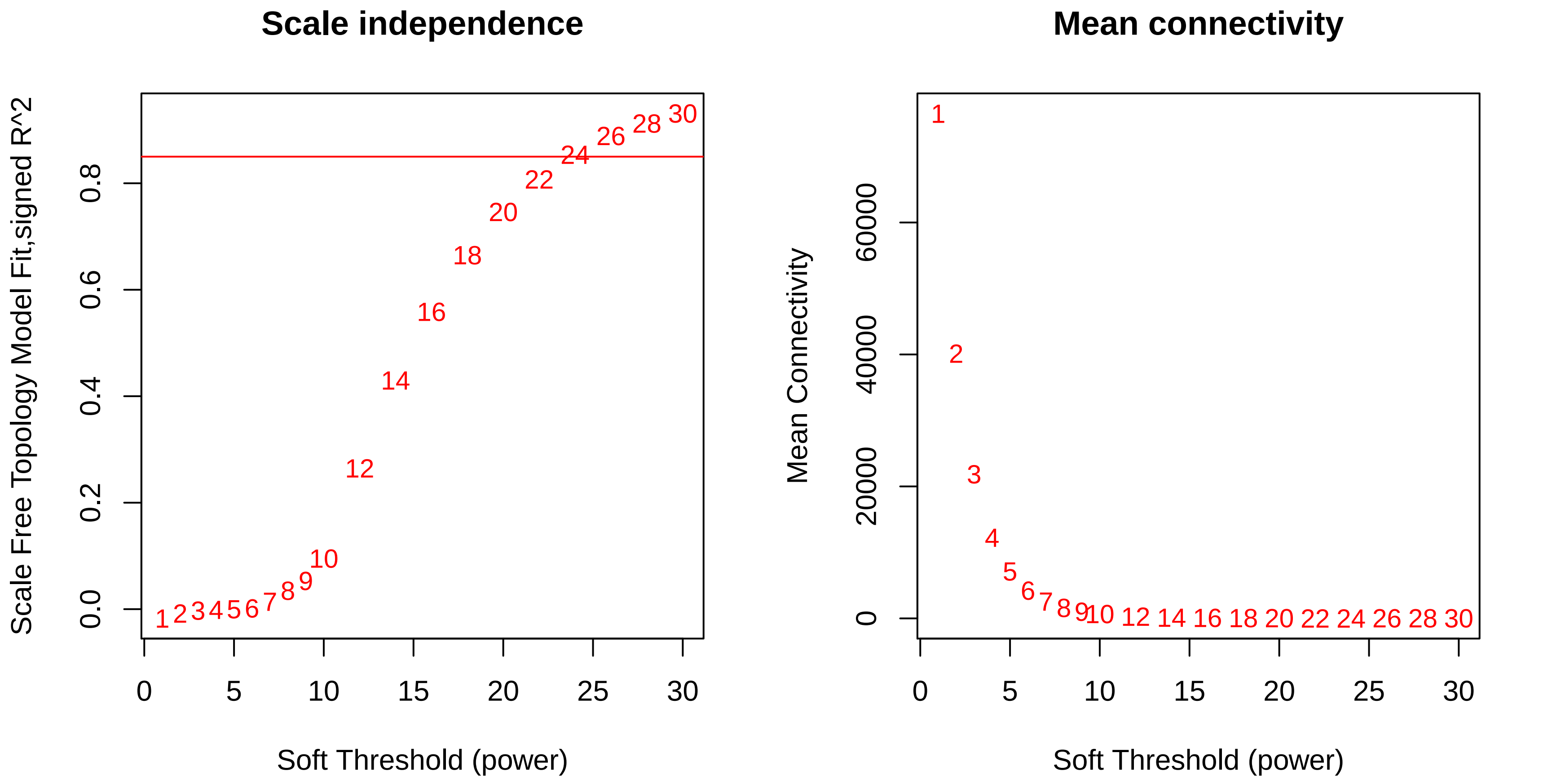

### Supplementary Figure 5

**Infant Hippocampus Module–trait Relationships**

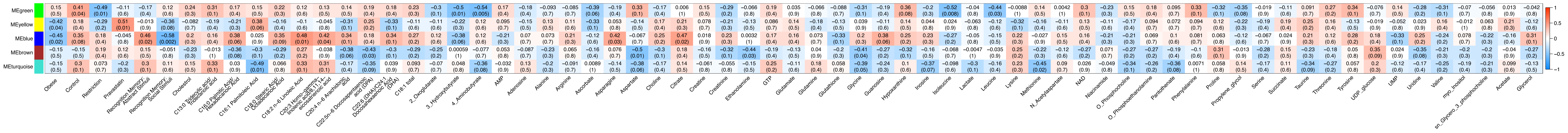

### Supplementary Figure 6

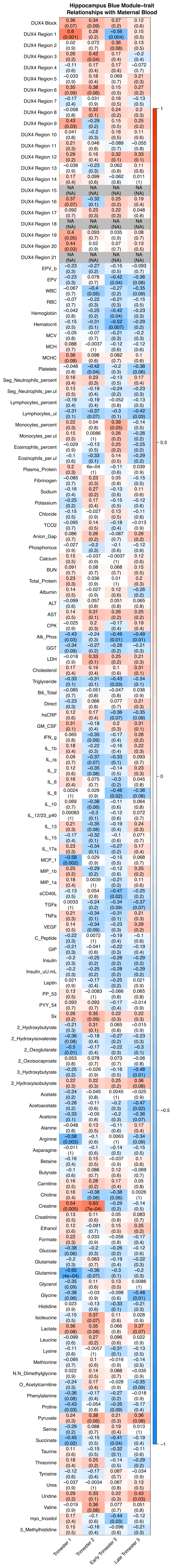
