## Supplementary Figure 3 for "Multi-omic brain and behavioral correlates of cell-free fetal DNA methylation in macaque maternal obesity models"

### mir663−trait Relationships with Infant Brain

| | Hippocampus | Hypothalamus | Prefrontal Cortex |
|---|---|---|---|
| Cholestrol | 0.077 (0.7) | 0.0089 (1) | −0.38 (0.06) |
| C13:0 Tridecylic acid/Tridecanoic acid | −0.18 (0.4) | −0.21 (0.3) | −0.15 (0.5) |
| C16:0 Palmitic Acid/Hexadecanoic Acid | −0.064 (0.8) | −0.12 (0.6) | −0.28 (0.2) |
| C16:1 Palmitoleic Acid | 0.036 (0.9) | −0.12 (0.6) | 0.34 (0.09) |
| C18:0 Stearic Acid/Octadecanoic Acid | −0.058 (0.8) | −0.012 (1) | −0.34 (0.1) |
| C18:2 n−6 Linoleic Acid | −0.21 (0.3) | −0.0032 (1) | −0.32 (0.1) |
| C20:3 Homo−gamma−linolenic acid/8,11,14−eicosatrienoic acid | −0.18 (0.4) | 0.0045 (1) | −0.42 (0.04) |
| C20:4 n−6 Arachidonic acid | −0.043 (0.8) | −0.13 (0.5) | −0.27 (0.2) |
| C22:5n−6 Docosapentaenoic acid (DPA) | 0.22 (0.3) | −0.12 (0.6) | 0.058 (0.8) |
| C22:6 (DHA)/C24:1 Docosahexaenoic Acid (DHA) | −0.01 (1) | −0.15 (0.5) | −0.32 (0.1) |
| C18:1 total | 0.012 (1) | 0.1 (0.6) | −0.32 (0.1) |
| 2_Oxoglutarate | 0.056 (0.8) | −0.048 (0.8) | −0.1 (0.6) |
| 3_Hydroxybutyrate | 0.17 (0.4) | 0.13 (0.5) | 0.16 (0.4) |
| 4_Aminobutyrate | −0.061 (0.8) | −0.0014 (1) | 0.25 (0.2) |
| AMP | 0.3 (0.1) | 0.062 (0.8) | −0.036 (0.9) |
| Adenosine | 0.089 (0.7) | 0.36 (0.08) | 0.5 (0.01) |
| Alanine | −0.12 (0.6) | −0.17 (0.4) | −0.024 (0.9) |
| Arginine | −0.16 (0.4) | 0.085 (0.7) | −0.25 (0.2) |
| Ascorbate | 0.096 (0.6) | 0.027 (0.9) | 0.022 (0.9) |
| Asparagine | −0.23 (0.3) | 0.32 (0.1) | −0.083 (0.7) |
| Aspartate | 0.071 (0.7) | 0.14 (0.5) | 0.074 (0.7) |
| Choline | −0.26 (0.2) | −0.27 (0.2) | −0.29 (0.2) |
| Citrate | −0.37 (0.07) | 0.16 (0.4) | −0.12 (0.6) |
| Creatine | 0.061 (0.8) | 0.046 (0.8) | 0.15 (0.5) |
| Creatinine | 0.0036 (1) | −0.2 (0.3) | 0.11 (0.6) |
| Ethanolamine | 0.14 (0.5) | 0.27 (0.2) | 0.21 (0.3) |
| GTP | 0.2 (0.3) | 0.45 (0.02) | 0.14 (0.5) |
| Glutamate | −0.042 (0.8) | 0.05 (0.8) | −0.083 (0.7) |
| Glutamine | −0.018 (0.9) | 0.056 (0.8) | 0.1 (0.6) |
| Glutathione | 0.35 (0.09) | 0.043 (0.8) | 0.066 (0.8) |
| Glycine | −0.078 (0.7) | −0.079 (0.7) | 0.16 (0.4) |
| Guanosine | −0.31 (0.1) | −0.023 (0.9) | 0.13 (0.5) |
| Hypoxanthine | −0.072 (0.7) | 0.11 (0.6) | −0.16 (0.5) |
| Inosine | −0.48 (0.02) | −0.36 (0.08) | −0.14 (0.5) |
| Isoleucine | 0.075 (0.7) | 0.063 (0.8) | 0.08 (0.7) |
| Lactate | −0.24 (0.2) | −0.16 (0.4) | −0.043 (0.8) |
| Leucine | −0.1 (0.6) | −0.067 (0.8) | −0.0043 (1) |
| Lysine | −0.31 (0.1) | 0.24 (0.3) | 0.2 (0.3) |
| Methionine | 0.058 (0.8) | 0.095 (0.7) | 0.2 (0.3) |
| N_Acetylaspartate | 0.058 (0.8) | 0.19 (0.4) | 0.13 (0.5) |
| NAD | 0.069 (0.7) | 0.48 (0.02) | −0.0019 (1) |
| Niacinamide | −0.3 (0.1) | −0.035 (0.9) | −0.089 (0.7) |
| O_Phosphocholine | −0.043 (0.8) | −0.027 (0.9) | 0.36 (0.07) |
| O_Phosphoethanolamine | 0.035 (0.9) | −0.059 (0.8) | 0.088 (0.7) |
| Pantothenate | −0.2 (0.3) | −0.13 (0.5) | −0.22 (0.3) |
| Phenylalanine | −0.02 (0.9) | −0.024 (0.9) | 0.042 (0.8) |
| Proline | −0.24 (0.2) | 0.34 (0.09) | −0.13 (0.5) |
| Propylene_glycol | −0.099 (0.6) | −0.13 (0.5) | 0.2 (0.3) |
| Serine | 0.035 (0.9) | −0.0028 (1) | 0.11 (0.6) |
| Succinate | −0.06 (0.8) | 0.17 (0.4) | 0.038 (0.9) |
| Taurine | 0.072 (0.7) | 0.055 (0.8) | 0.23 (0.3) |
| Threonine | −0.17 (0.4) | −0.35 (0.09) | 0.2 (0.3) |
| Tyrosine | −0.22 (0.3) | −0.21 (0.3) | −0.15 (0.5) |
| UDP_glucose | −0.066 (0.8) | 0.18 (0.4) | −0.0068 (1) |
| UMP | 0.093 (0.7) | 0.0098 (1) | −0.18 (0.4) |
| Uridine | −0.3 (0.1) | −0.006 (1) | 0.12 (0.6) |
| Valine | 0.056 (0.8) | 0.059 (0.8) | 0.18 (0.4) |
| myo_Inositol | −0.048 (0.8) | 0.14 (0.5) | 0.03 (0.9) |
| sn_Glycero_3_phosphocholine | −0.061 (0.8) | −0.071 (0.7) | 0.21 (0.3) |
| Acetate | −0.25 (0.2) | −0.0037 (1) | −0.12 (0.6) |
| Glycerol | −0.21 (0.3) | −0.26 (0.2) | −0.035 (0.9) |

Color scale: 1, 0.5, 0, −0.5, −1
